# Hypoxia-Induced Ethanol Production in Arabidopsis Root Apex Cells Affects Endocytic Vesicle Recycling and F-Actin Organisation

**DOI:** 10.1101/2022.01.05.475021

**Authors:** Tomoko Kagenishi, František Baluška, Ken Yokawa

## Abstract

Ethanol (EtOH) is a short-chain alcohol that is abundant in nature. EtOH is endogenously produced by plants under hypoxic conditions, and exogenously applied EtOH improves plant stress tolerance at low concentrations (<1%). However, no direct observations have shown how EtOH affects cellular events in plants. In intact *Arabidopsis* roots, 0.1% EtOH promoted reactive oxygen species production in root apex cells. EtOH also accelerated exocytic vesicle recycling and altered F-actin organisation, both of which are closely related to cell membrane properties. In addition to exogenous EtOH application, hypoxic treatment resulted in EtOH production in roots and degradation of the cross-wall actin cytoskeleton in root epidermal cells. We conclude that hypoxia-induced EtOH production affects endocytic vesicle recycling and associated signalling pathways.

## Introduction

Ethanol (EtOH) is a small volatile molecule that is abundant in nature. EtOH is produced through multiple chemical and biochemical reactions and in particular during yeast-mediated fermentation. Extensive consumption of EtOH produces a drunken state in animals and humans due to the molecular interactions with certain components of the nervous system. Plants are also susceptible to the effects of EtOH. High concentrations of EtOH can damage plants; however, low concentrations have been shown to moderate plant stress responses. For example, incubating cucumber seedlings in EtOH vapour (<3%) induced chilling tolerance [1]. Similarly, the application of 0.3% ethanol enhanced the salinity tolerance of Arabidopsis and rice roots [2,3]. Although EtOH treatments also promoted the expression of heat shock proteins in soybean, no improvements in heat tolerance were observed [4].

Some plants can also produce EtOH under hypoxic and anoxic conditions. During this process, pyruvate is converted into acetaldehyde by pyruvate decarboxylase, then acetaldehyde is converted into EtOH by alcohol dehydrogenase (ADH) [5–10]. This reversible enzymatic reaction has been observed in yeast, bacteria, *Chlamydomonas* [11] and plants [12] under low oxygen conditions. Arabidopsis roots constitutively express the *ADH* gene, and Chung *et al*. [13] demonstrated that ADH is strongly upregulated when roots are exposed to low oxygen conditions upon penetrating solid-agar media. Hypoxic and anoxic conditions are sensed by roots and subsequently induce ADH expression in shoots, which suggests that EtOH plays a role in communication between shoots and roots [13]. Furthermore, low temperatures induce ADH expression and EtOH biogenesis in shoots and roots of rice seedlings, and EtOH treatments improve the cold tolerance of rice seedlings [14]. Together these observations suggest that in plants, endogenous EtOH is a stress-induced molecule that facilitates physiological adaptations to changes in environmental conditions.

Although numerous studies have investigated the role of EtOH in plants, the initial events and cellular components that mediate plant responses to EtOH remain poorly understood. In previous studies, EtOH was reported to fluidise *Escherichia coli* plasma membranes [15]; molecular dynamics simulations showed that EtOH interacts with membrane lipids [16]. The plasma membrane is a highly active and dynamic structure in all organisms, and the elaborate control of membrane homeostasis is crucial to many cellular events. In human astrocytes, 0.15% EtOH treatments disrupted the actin cytoskeleton structures [17]. Thus, it is accepted that even small membrane disturbances can have substantial impacts on membrane proteins, ion channels and endo/exocytic vesicle recycling [18]. To our knowledge, the present study is the first to have monitored the membrane-associated effects of EtOH in intact Arabidopsis root cells. Under hypoxic conditions, endogenous EtOH production was observed in the root tissue. We used confocal microscopy to demonstrate that EtOH inhibits Arabidopsis root growth and interrupts vesicle recycling. In addition, we observed reactive oxygen species (ROS) accumulation in the root apex regions and perturbations of F-actin polarisation under both EtOH and hypoxic treatments.

## Results and Discussion

### Ethanol production under hypoxic conditions in Arabidopsis roots

Many plants produce considerable quantities of ethanol under anoxic and hypoxic growth conditions [5–7,9,10]. For example, Kato-Noguchi (2000) demonstrated that EtOH levels in lettuce roots increased following hypoxic treatments [8]. Here we first used spectrophotometry to determine whether Arabidopsis roots produce EtOH under low oxygen conditions. As shown in Fig. 1, the endogenous EtOH concentrations were significantly increased after incubation in anoxic conditions (nitrogen gas) for 24 h. The range of EtOH concentrations detected here (μmol g^-1^ FW order) was consistent with a previous report [10]. EtOH production was also observed in our untreated controls, presumably due to the penetration of horizontally-incubated roots into the watery solid medium, as described previously [13]. In a recent study, Eysholdt-Derzsó and Sauter revealed that hypoxic conditions activate the group VII ethylene response factor (ERFVII), resulting in slanted root growth, possibly as a mechanism to avoid low oxygen conditions [19]. Changes in auxin distribution and signalling in root tips have also been reported under hypoxic conditions [19]. Next, we monitored changes in the root growth and gravitropism response following exogenous application of EtOH.

**Figure 1.**
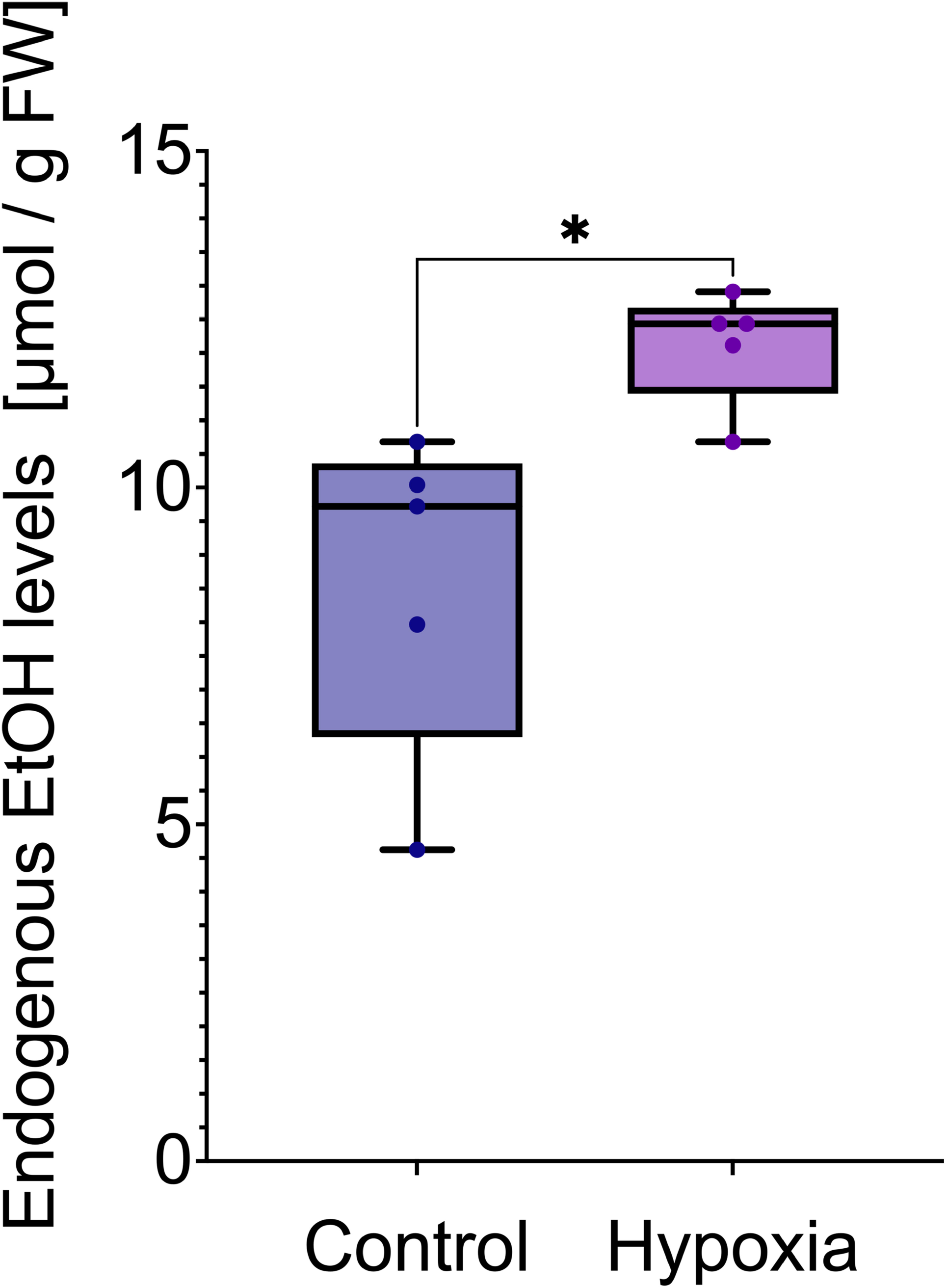
Hypoxia-induced EtOH production in Arabidopsis root tissues. Ethanol contents were biochemically determined using a spectrophotometer based on the amount of NADH metabolised by alcohol dehydrogenase and acetaldehyde dehydrogenase. Seedlings were treated with pure nitrogen gas for 24 h to induce hypoxia. Error bars indicate standard deviations of the mean from five independent experiments. Each sample contained approximately 40 mg of root tissue (~100 roots). Significant differences were identified using Student’s t-test; \**P* < 0.05.

### Inhibition of root growth by EtOH

We monitored Arabidopsis root growth in 0.5 × Murashige-Skoog (MS) medium containing 0, 0.01%, 0.1% and 1% EtOH. The experimental EtOH concentrations did not affect seed germination (data not shown). However, as shown in Fig. 2A, 0.01%, 0.1% and 1% EtOH significantly inhibited root growth; the inhibition effect was strongest in the 1% EtOH treatment. Arcara and Ronchi [20] reported that treating onion (*Alium cepa* L.) with 0.4% EtOH delayed the mitotic cycle, but did not completely stop mitosis. The root growth inhibition observed here is likely the result of a similar response to EtOH in *Arabidopsis*. In contrast, root gravitropism was not affected by EtOH, even after 24 h in 1% EtOH (Fig. 2B). The DR5 auxin reporter expression was not affected, and auxin redistribution after 6 h gravistimulation in the presence of 0.1% or 1% EtOH was not significantly different from the control (Fig. 2C). Thus, EtOH at the concentrations tested did not affect the events required for graviresponse such as auxin redistribution.

**Figure 2.**
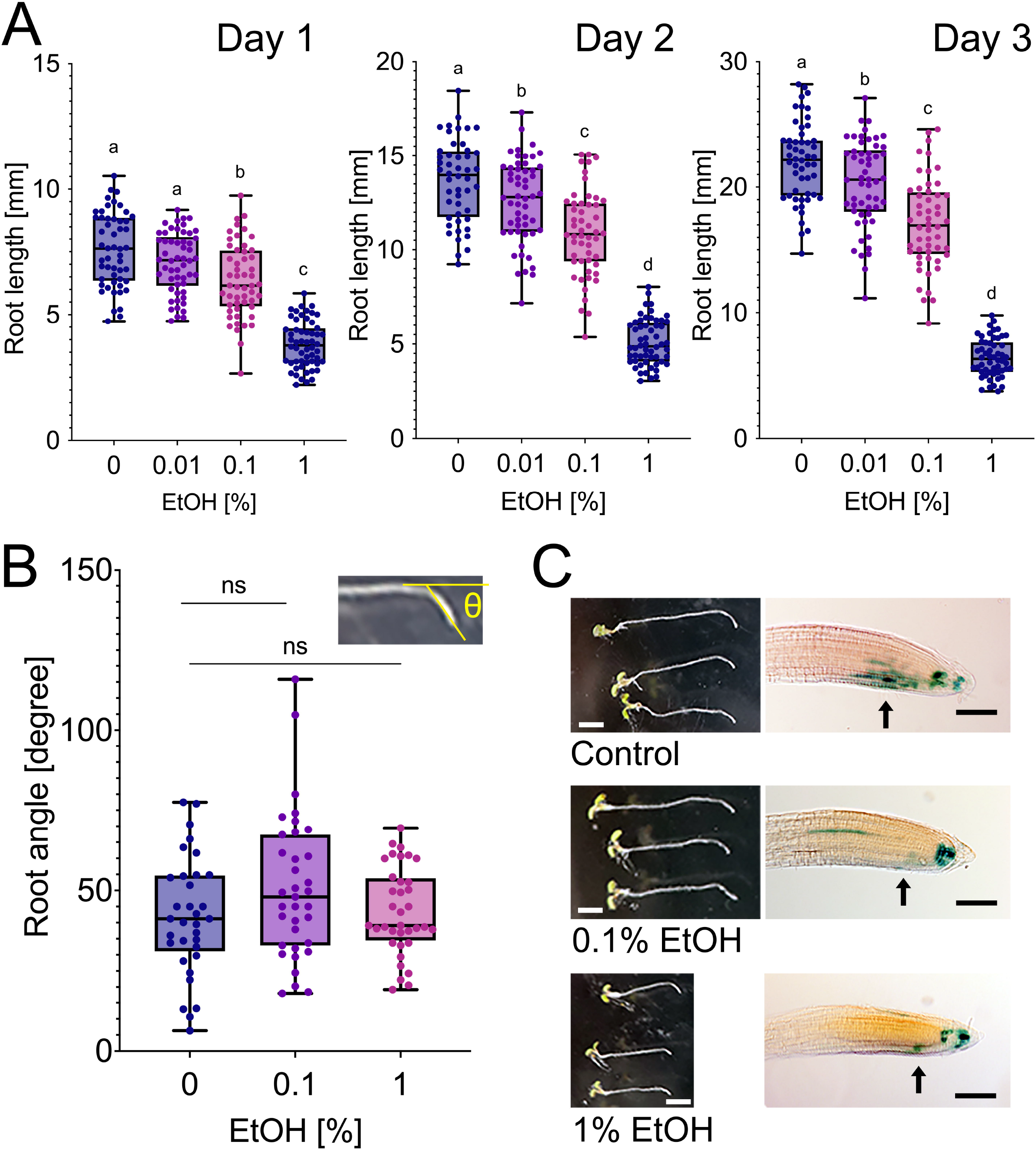
Root lengths are affected by EtOH treatments but not gravitropism. (A) Lengths of roots grown in the presence of 0%, 0.01%, 0.1% or 1% EtOH for 3 days (n = 52-56 with three biological replicates). Different letters indicate significant differences (Tukey’s HSD test, *P* < 0.05). (B) Angles of root gravitropic curvature in the presence of EtOH. The x-axis shows the treatment time after gravistimulation started. There were no significant differences in each EtOH treatment time. Error bars indicate standard deviations of the mean (n = 33-36 with three biological replicates). (C) Gravitropic curvature in the presence of EtOH. The left column shows root gravitropism under three different EtOH concentrations. Scale bar = 5 mm. The right column shows the expression of auxin reporter mutant DR5::GUS. Scale bar = 100 μm.

### EtOH induces ROS accumulation in root apex

ROS in root apex cells play a crucial role in root development and growth [21]. The accumulation of ROS has been observed in EtOH-treated yeast [22]. In hepatocytes, ROS production was induced by EtOH concentrations as low as 1 mM (~0.006%) [23]. To determine whether ROS homeostasis is altered in plant roots in the presence of EtOH, we visualised superoxide localisation in Arabidopsis root apices using nitroblue tetrazolium salt (NBT) staining (Fig. 3). Superoxide production was promoted by 0.1% and 1% EtOH, as demonstrated by the accumulation of superoxide in root apex regions. According to a previous report [21], the balance of superoxide and hydrogen peroxide is strictly regulated in these regions. Moreover, hydrogen peroxide promotes cell elongation, whereas superoxide promotes cell proliferation. Hence, superoxide accumulation under EtOH treatments is consistent with the inhibition of root growth but not with root responses to gravity. Nguyen *et al*. [2] recently showed that plant salt tolerance is improved by treatments with exogenous EtOH, reflecting the induction of ROS detoxifying enzymes such as peroxidases. Here we analysed the gene expression of transcription factor ZAT10, which provides an ROS signalling-dependent stress response [24]. After treatment with 1% EtOH, the ZAT10 expression level was about four-fold higher in the control. Hence, EtOH-mediated ROS production may regulate the transcription of genes, resulting in the regulation of antioxidant enzymes. However, further studies are required to determine the mechanisms and sources of ROS production.

**Figure 3.**
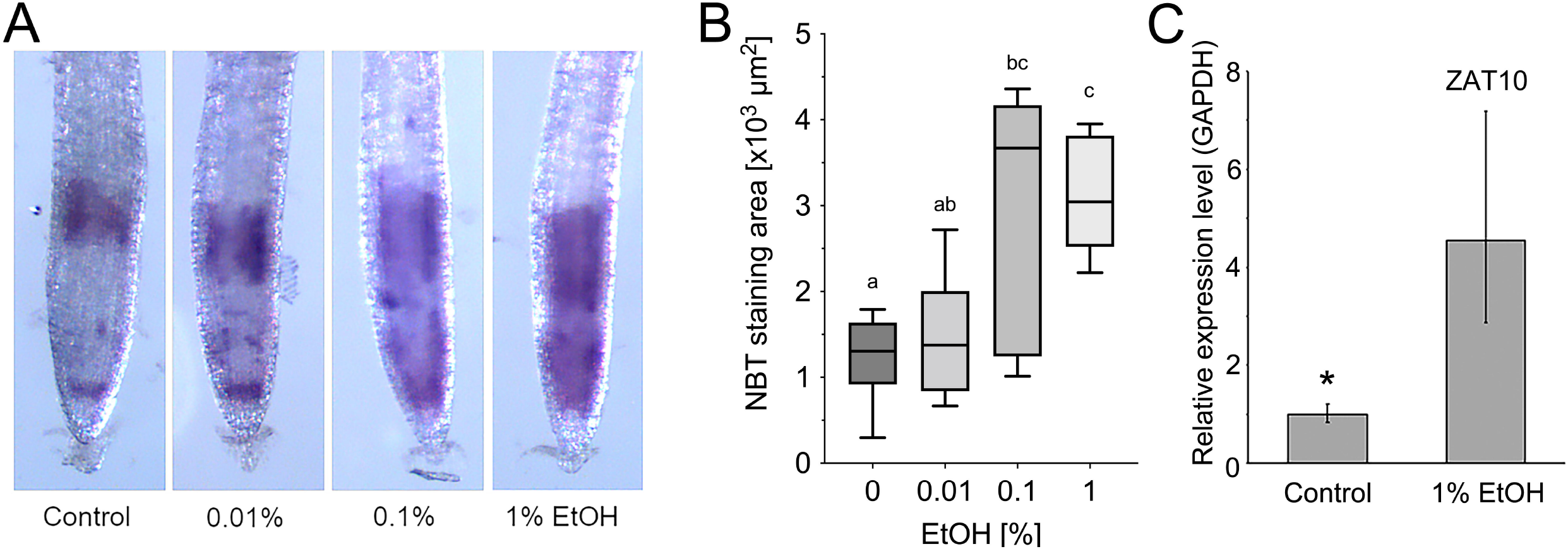
Ethanol induces the production of superoxide in Arabidopsis root apices. (A) Seedlings were treated with EtOH for 1 h and then stained with nitroblue tetrazolium salt (NBT) solution to visualise superoxide distribution. (B) NBT-stained areas were quantified. Error bars indicate standard deviations from the mean (n = 5-6 with three biological replicates). Different letters indicate significant differences (Tukey’s HSD test, *P* < 0.05). (C) Real-time PCR analysis for ROS signalling-dependent transcription factor, ZAT10; expression was analysed using the ΔΔCt method with GAPDH3 as the reference gene.

### EtOH unbalances between endocytic and exocytic vesicle recycling pathways

Because EtOH directly interacts with membrane structures, we assessed endocytic vesicle recycling in the presence of EtOH in root epidermal cells. We used brefeldin A (BFA) to monitor membrane recycling in Arabidopsis. BFA prevents the formation of exocytic vesicles but not endocytic vesicles [25]; in particular, the area of BFA compartments is indicative of the rate of membrane recycling through the endocytic pathway. As shown in Figs 4A and B, treatments with 0.1% EtOH did not affect the BFA-induced compartment area. However, 30 min of BFA treatment after 1% EtOH pre-treatment inhibited the endocytic vesicular pathways that reflect small BFA-induced compartments in Arabidopsis seedlings (Figs 4A and B). BFA does not modify dopamine receptor localisation, indicating that endosomal recycling is accomplished through BFA-independent pathways [26]. By washing the BFA from the EtOH-treated roots, the retarded exocytic pathway traffic is released again and the speed of exocytosis can be monitored. Our BFA washing-out experiments revealed that 0.1% or 1% EtOH treatments sped up the outward exocytic pathway 30 min after BFA removal (Figs 4C and D). Taken together, these data suggest that EtOH modifies BFA-sensitive signalling pathways in plants. Inward endocytic pathway was inhibited and outward exocytic one was accelerated. Hypoxia-induced root slanting occurs via the alteration of auxin distribution [19]; however, our data show that exogenous EtOH does not affect DR5 expression and root gravitropic responses (Figs 2B and C). These findings imply that the acceleration of exocytic vesicle recycling does not affect gravitropic responses and auxin distribution for up to 24 h of EtOH treatment.

**Figure 4.**
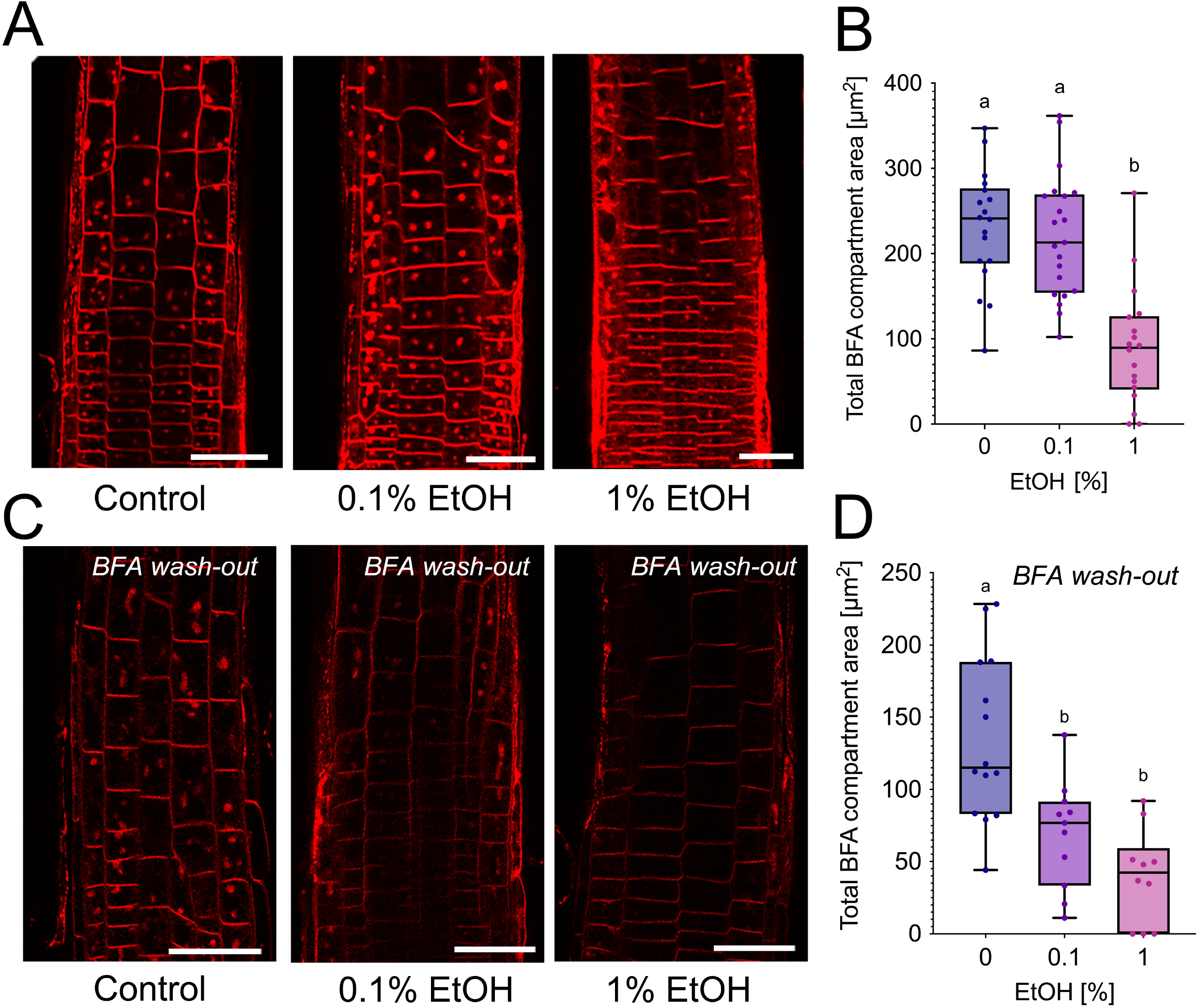
BFA-induced compartments in root apex transition zone cells exposed to EtOH. (A) Confocal images of BFA-induced compartments in the root epidermis of the transition zone after EtOH treatments. Five-day-old seedlings were stained with 4-μM FM4-64 for 10 min, incubated in 0% or 1% EtOH in 0.5 × MS for 1 h and then soaked in 35-μM BFA in 0.5 × MS for 30 min. Scale bar = 50 μm. (B) The total area of BFA-induced compartments in root transition zone epidermis cells was measured using ImageJ (n = 18-21 with three biological replicates). Error bars indicate standard deviations from the mean. (C) Recovery from BFA-mediated inhibition of vesicle recycling following washing. After FM4-64 staining, seedlings were soaked in 35-μM BFA in 0.5 × MS for 30 min and then incubated in 0% or 1% EtOH in 0.5 × MS for 1 h. Scale bar = 50 μm. (D) Reductions in BFA compartment sizes after recovery. The total area of BFA-induced compartments in root transition zone epidermis cells was measured using ImageJ (n = 10-14 with three biological replicates). Error bars indicate standard deviations from the mean; ns, not significant; different letters indicate significant differences according to Tukey’s HSD test, *P* < 0.05.

### Dispersion of F-actin bundles in root apex region cross-walls

F-actin acts as a scaffold system for moving vesicles in the root transition zone [25]. The influence of EtOH on F-actin localisation in root epidermal cells was observed in GFP-FABD2 transgenic *Arabidopsis* plants [27]. As shown in Fig. 5A, F-actin alignment at the cross-wall (white arrow) disappeared in the presence of 0.1% and 1% EtOH. The effects of EtOH on F-actin have been reported in other organisms. For example, F-actin in budding yeast was dispersed following induction by 6% EtOH [28]. EtOH treatments at 0.15% concentration clearly disrupted actin cytoskeleton structures in astrocytes [17]. In mouse cells, the actin cytoskeleton acts as both the target and effector of the EtOH effects on synapses, as indicated by disturbances in N-methyl-D-aspartate receptor signalling [29–31]. Furthermore, abundant F-actin and active endocytic vesicle recycling were previously observed in root apex transition regions [32–34]. Thus, the depletion of cross-wall localised F-actin in EtOH-treated root apex cells may disturb vesicle recycling, root growth and polarity, as well as influencing the general cell physiology.

**Figure 5.**
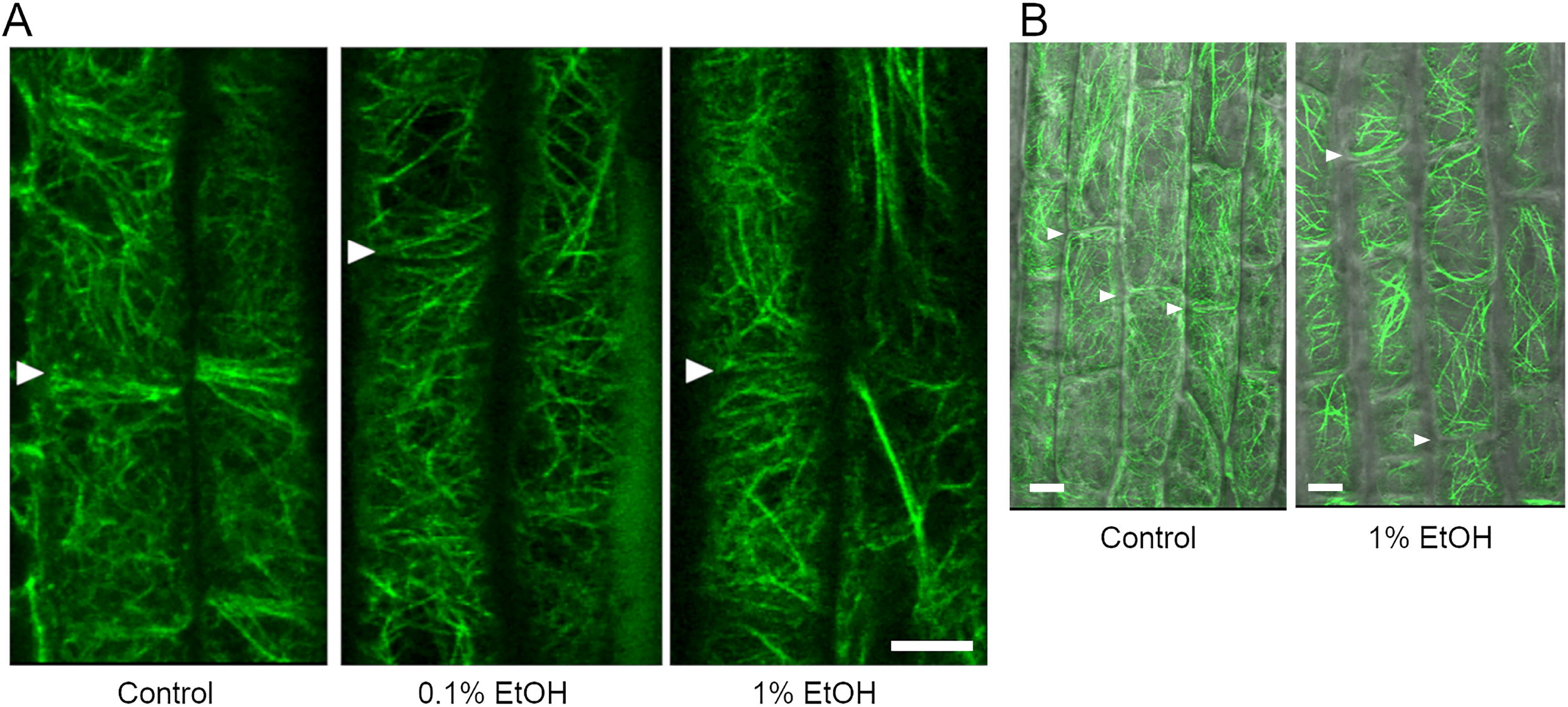
Visualisation of F-actin in epidermal cells in the root elongation zone. (A) Five-day-old light-grown GFP-FABD2 seedlings were incubated in 0%, 0.1% or 1% EtOH in 0.5 × MS solid media for 1 h. The cross-wall position under each condition is indicated with an arrow. Five replicates produced similar results and representative examples are presented for each condition. (B) Enlarged confocal image corresponding to Figure 5A. The cross-wall position under each condition is indicated with an arrow.

### Depletion and reorganisation of F-actin under hypoxia

To clarify whether hypoxic conditions change F-actin organisation without exogenous EtOH application, we monitored GFP-FADB2 root transition zone cells under hypoxia for 1, 3, 24 h. F-actin depletion was observed after 1-h hypoxic treatment and the signal disappeared from the cross-walls in the epidermal cells (Fig. 6A). The timing of GFP signal loss from the cross-walls was the same under the 1-h hypoxic and 0.1% EtOH treatments (Fig. 5A). This finding suggests that endogenous EtOH production under hypoxia may be initiated as soon as roots are exposed to low oxygen conditions. The accumulated F-actin bundle moved to the tonoplast surface under hypoxic conditions (Fig. 6B). Importantly, F-actin recovery following the 24-h hypoxia treatment was observed after just 1 h (Fig. 6C), suggesting that hypoxia-induced F-actin disruption may be mediated by endogenous EtOH production. Interestingly, the root apex cells in the transition zone show the most rapid and extensive F-actin redistribution [35]. This zone also contains cells with the highest demands for oxygen uptake [36,37], rates of endocytic vesicle recycling and auxin fluxes [39–40].

**Figure 6.**
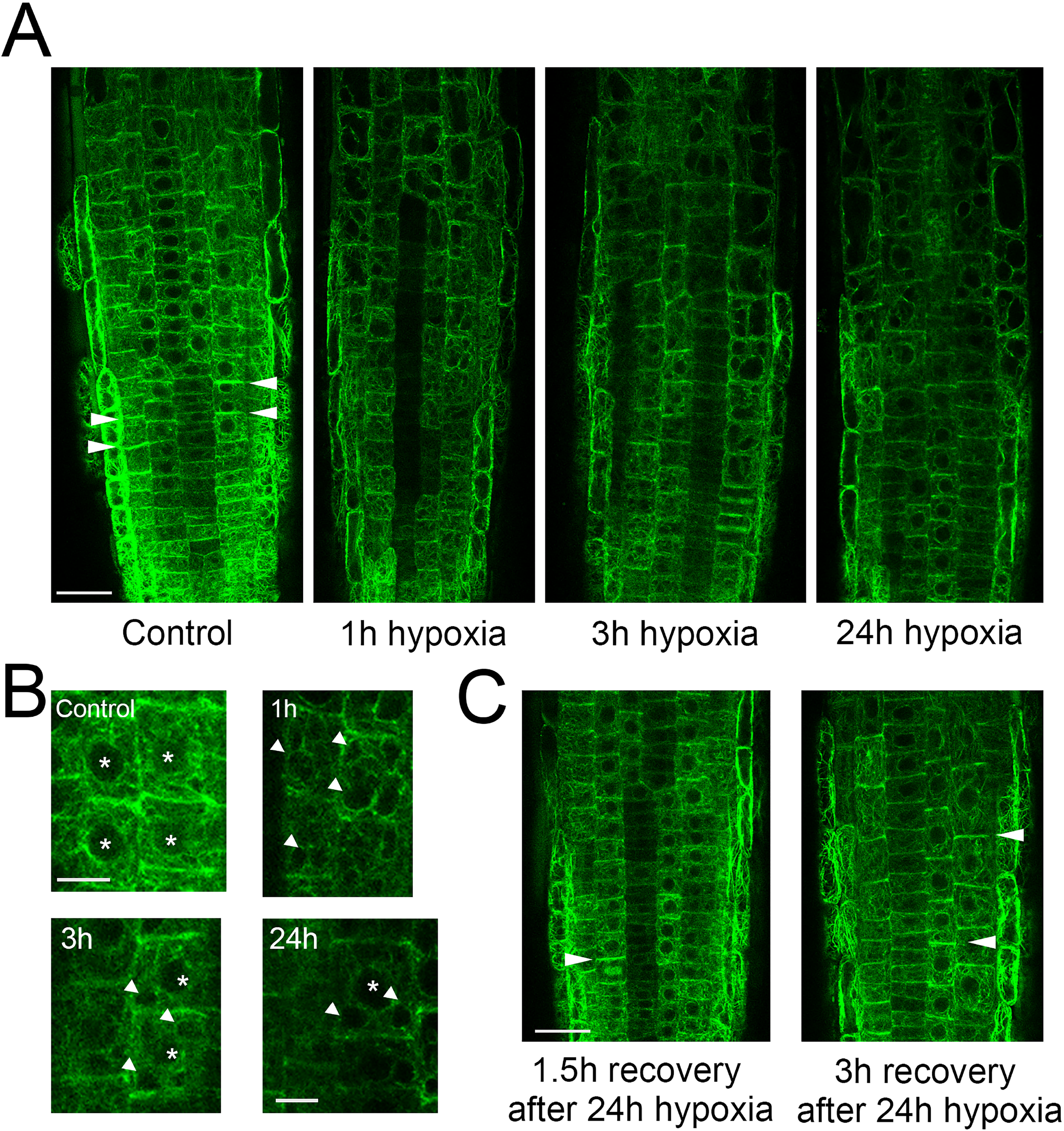
Hypoxia-induced depletion of F-actin in Arabidopsis GFP-FABD2 root apex. (A) Hypoxia-induced the depolymerisation of F-actin after just 1 hour. Both cytosolic and cross-wall F-actin were depleted. The position of the cross-wall is indicated with an arrow. Scale bar = 25μm. (B) The enlarged images from (A). An asterisk and an arrow indicate the position of the nucleus and F-actin accumulation on the tonoplast membrane in the treated root cells, respectively. Scale bar = 10μm. (C) F-actin recovery from hypoxic conditions. Scale bar = 25μm. The position of the cross-wall is indicated with an arrow. Representative images from six roots are shown.

### Conclusions

We conclude that EtOH produced under hypoxic conditions directly affects the physical properties of plasma membranes, contributing to ROS homeostasis, vesicle recycling and F-actin organisation. These EtOH-responsive processes help roots to rapidly and effectively adapt to hypoxic environments. The molecular details and spatio-temporal relationships between EtOH-modified membranes, endocytic vesicle recycling and F-actin organisation will be investigated in future studies.

## Materials and Methods

### Plant growth conditions and exogenous EtOH treatments

*Arabidopsis thaliana* seeds were sterilised in 2% NaClO containing 0.1% Triton X-100 for five minutes. Seeds were then washed with distilled water four times and planted in Petri dishes on 0.5 × MS basal salt medium containing 1% (w/v) sucrose (pH 5.8 with KOH) solidified with 0.4% (w/v) phytagel containing 0, 0.01, 0.1 or 1% EtOH (VWR chemicals, France). Approximately 20 seedling were placed on EtOH-containing plates of each concentration and three times biological replicates were conducted. Ecotype Col-0 was used for the wild type, the 35S: GFP-FABD2 transgenic line was used for F-actin visualisation [27] and DR5::GUS was used for visualising the localisation of auxin during gravitropism. The MS medium was autoclaved and sufficiently cooled before adding the EtOH to prevent evaporation. The Petri dishes were incubated at 4°C for one day, then incubated vertically at 23°C–25°C under a 16-h light/8-h dark cycle. Light was provided at an intensity of 120 μmol/m^2^/s. Three days after germination, images of roots were taken using an EOS Kiss X7 camera (Canon, Tokyo) and root lengths were measured using ImageJ software (ver. 1.43u for Mac OSX, http://imagej.nih.gov/ij/). The three-day-old seedlings were transferred to EtOH containing 0.5 × MS solid media (0%, 0.1% or 1%) and rotated 90 degrees to observe root gravitropism. The seedlings were then incubated in the dark for 24 h. Images to assess root curvature were captured and measured as described above.

### Hypoxic treatments and endogenous EtOH production measurement

For the hypoxia treatments, seedlings growing on 0.5 × MS solid medium in Petri dishes (without lids) were placed in a gas-tight chamber five days after germination. To expel oxygen, pure nitrogen gas was infused into the closed chamber through small holes for 5 min. The holes were then tightly sealed and the plates in the chamber were incubated horizontally at 23°C–25°C for 24 h in the dark to prevent photosynthesis. Control seedlings were prepared without nitrogen gas treatments. After the hypoxic treatments, roots were cut from the seedlings and weighed. The root samples were then ground in liquid nitrogen using a pestle and mortar. Subsequently, 300-μL aliquots of distilled water were added and mixed with the ground powder. The mixture was centrifuged in 1.5 mL tubes at 15,000 rpm (21,386 × g) for 1 min at 1°C and the supernatant was collected. The EtOH content was then determined spectrophotometrically using a commercial kit (Ethanol, R-Biopharm AG, Germany) according to the manufacturer’s instructions. Briefly, the kit contained nicotinamide-adenine dinucleotide (NAD), alcohol dehydrogenase and aldehyde dehydrogenase. In spectrophotometric reactions, alcohol dehydrogenase oxidises EtOH to acetaldehyde in the presence of NAD, generating NADH. Aldehyde dehydrogenase also oxidises acetaldehyde to acetic acid and generates NADH in the presence of NAD. Hence, NADH absorbance at 334, 340 and 365 nm was determined using a spectrophotometer (U-5100, Hitachi, Japan) and stoichiometry was used to calculate the root extract EtOH concentrations using the extinction coefficient of NADH.

### Histochemical root staining

For the GUS staining, Petri dishes with five-day-old DR5::GUS seedlings were rotated 90 degrees and placed in the dark for 6 h. After gravistimulation, images of the seedlings were captured using an EOS Kiss X7 camera (Canon, Tokyo) and fixed with 90% cold acetone for 15 min. The acetone was discarded and the seedlings were rinsed with GUS buffer three times. The GUS buffer contained 50-mM potassium phosphate buffer (pH = 7.2), 0.5-mM potassium ferricyanide and 0.5-mM potassium ferrocyanide. Next, the seedlings were stained with 1-mM X-Gluc/DMF dissolved in GUS buffer in a vacuum chamber. The stained samples were incubated overnight at 37°C and rinsed with 70% EtOH before observation.

To detect superoxide in root apices grown in various concentrations of EtOH, five-day-old seedlings were incubated in EtOH solution for 1 h before staining in a solution containing 300-μM NBT (Fluka, Germany) dissolved in 0.1-M Tris-HCl, 0.1-M NaCl and 0.05-M MgCl_2_ (pH = 9.5) for 5 min. Seedlings were then observed and imaged under a light stereomicroscope. The NBT-stained areas in the binarised images with identical threshold were measured using ImageJ.

### Quantification of gene expression in EtOH-treated roots

After the EtOH treatments, roots were cut from the seedlings and weighed. The root samples were frozen with liquid nitrogen and ground to a powder using a pestle and mortar. Total RNA was purified using an ISOSPIN Plant RNA kit (NIPPON GENE, Japan) following the manufacturer’s manual. Single-strand cDNA was also obtained using SuperScript IV Reverse Transcriptase (Thermo-Fischer, Japan). DNA amplification was conducted using a real-time PCR device, LightCycler Nano (Roche, Switzerland) with the following primers: ZAT10-forward, AGGCTCTTACATCACCAAGATTAG; ZAT10-reverse, TACACTTGTAGCTCAACTTCTCCA; GAPDH3-forward, TTGGTGACAACAGGTCAAGCA; GAPDH3-reverse, AAACTTGTCGCTCAATGCAATC. The relative gene expression was calculated using the ΔΔCt method with GAPDH3 (glyceraldehyde-3-phosphate dehydrogenase) as the reference gene.

### Confocal observations of endo/exocytic vesicle recycling

Five-day-old seedlings grown under the conditions described above were stained with the membrane-staining fluorescence probe FM4-64 at 4 μM (Sigma, Germany) for 10 min. The seedlings were then treated with 0, 0.1 or 1% (v/v) EtOH in 0.5 × MS for 1 h, followed by incubation in 35-μM BFA (Sigma, Germany) for 30 min. Confocal images of the seedlings were taken using a confocal laser microscope (Fluoview FV1000, Olympus, Tokyo, Japan) and the effects of EtOH on the endocytosis rates were compared. To measure exocytosis, we monitored seedling recovery from BFA-inhibited recycling. The seedlings were stained with 4-μM FM4-64 and then soaked in 35-μM BFA for 30 min. After washing out the BFA, the seedlings were incubated in EtOH at the stated concentrations for 1 h before confocal imaging. The FM4-64 dye was excited at 515 nm using an Argon laser and fluorescence emissions of FM4-64 were measured at 630–700 nm. To quantify the BFA compartment size in the root transition zone, confocal images of the epidermal cells were trimmed to a 50 μm square (10–15 cells). The plasma membrane red fluorescence was removed and the BFA compartments were extracted by adjusting the threshold in ImageJ. The total BFA compartment area (~30–50 particles) was calculated for each image and averaged.

### F-actin bundle visualisation

Transgenic *Arabidopsis* GFP-FABD2 (actin-binding domain 2) seedlings [27] were grown under the conditions described above. After five days, the seedlings were prepared for confocal microscopy analysis by incubation in 0.5 × MS containing EtOH for 2 h. GFP-FABD2 was excited at 488 nm using an Argon laser and the emissions were measured at 500–550 nm.

### Statistical analysis

All numerical data were analysed using Student’s t-test and Tukey’s honestly significant difference (HSD) using Prism 9 for macOS (ver. 9.3.1 (350), December 7, 2021). Differences were considered significant when *p* < 0.05.

## Acknowledgements

We thank to Dr. Yutaka Kodama (University of Utsunomiya) for his critical reading and suggestion. We also thank to Dr. Yoshihito Kohari (Kitami Institute of Technology) for his technical advice for EtOH quantification. This work was supported by JSPS KAKENHI, Grant-in-Aid for No. 18K14726 (T.K.) and in part by Leading Initiative for Excellent Young Researchers of MEXT, Japan (K.Y.). The authors would like to thank Enago (www.enago.jp) for the English language review.

## Author Contribution Statement

K.Y. and F.B coordinated the project. T.K. and K.Y. designed the experiments. T.K. performed the microscopy and biochemical analysis. K.Y. analysed the data and performed all calculations. All authors wrote the manuscript and agreed with the final manuscript.

